## Supplementary figures and images for "Subregional activity in the dentate gyrus is amplified during elevated cognitive demands" (from another version of this preprint: version 4, 2025-12-30)

### Supp Fig 1

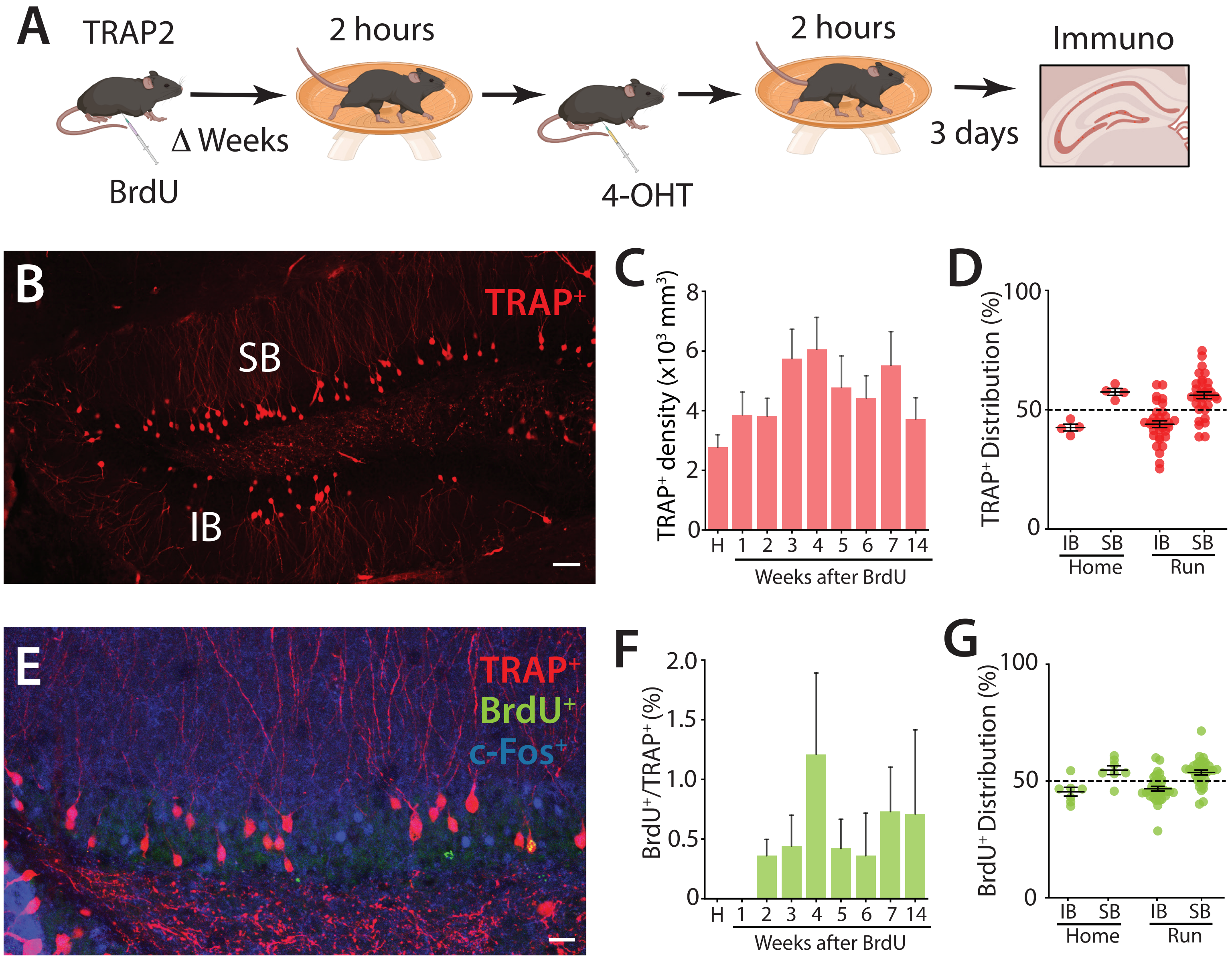

### Supp Fig 2

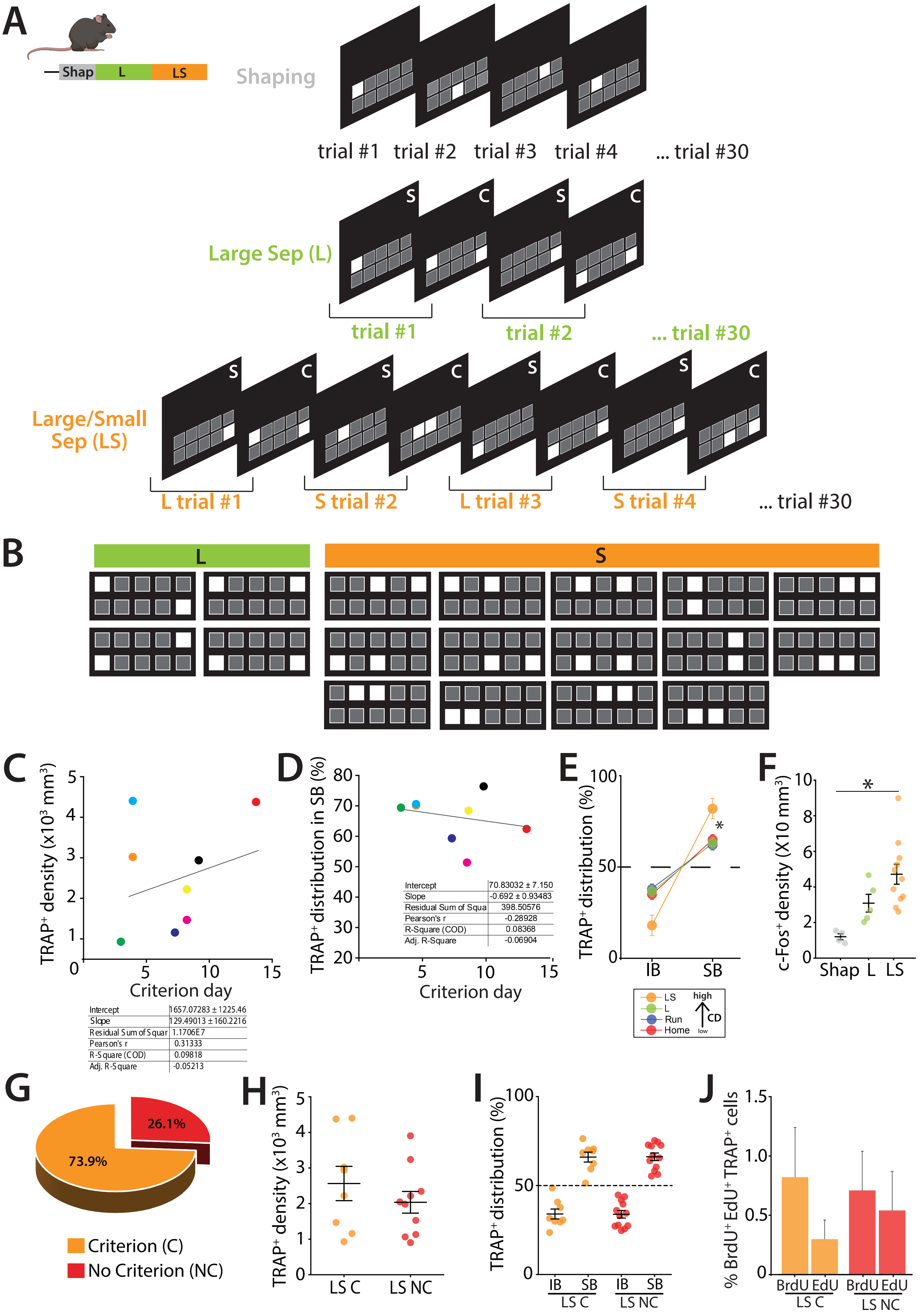

### Supp Fig 3

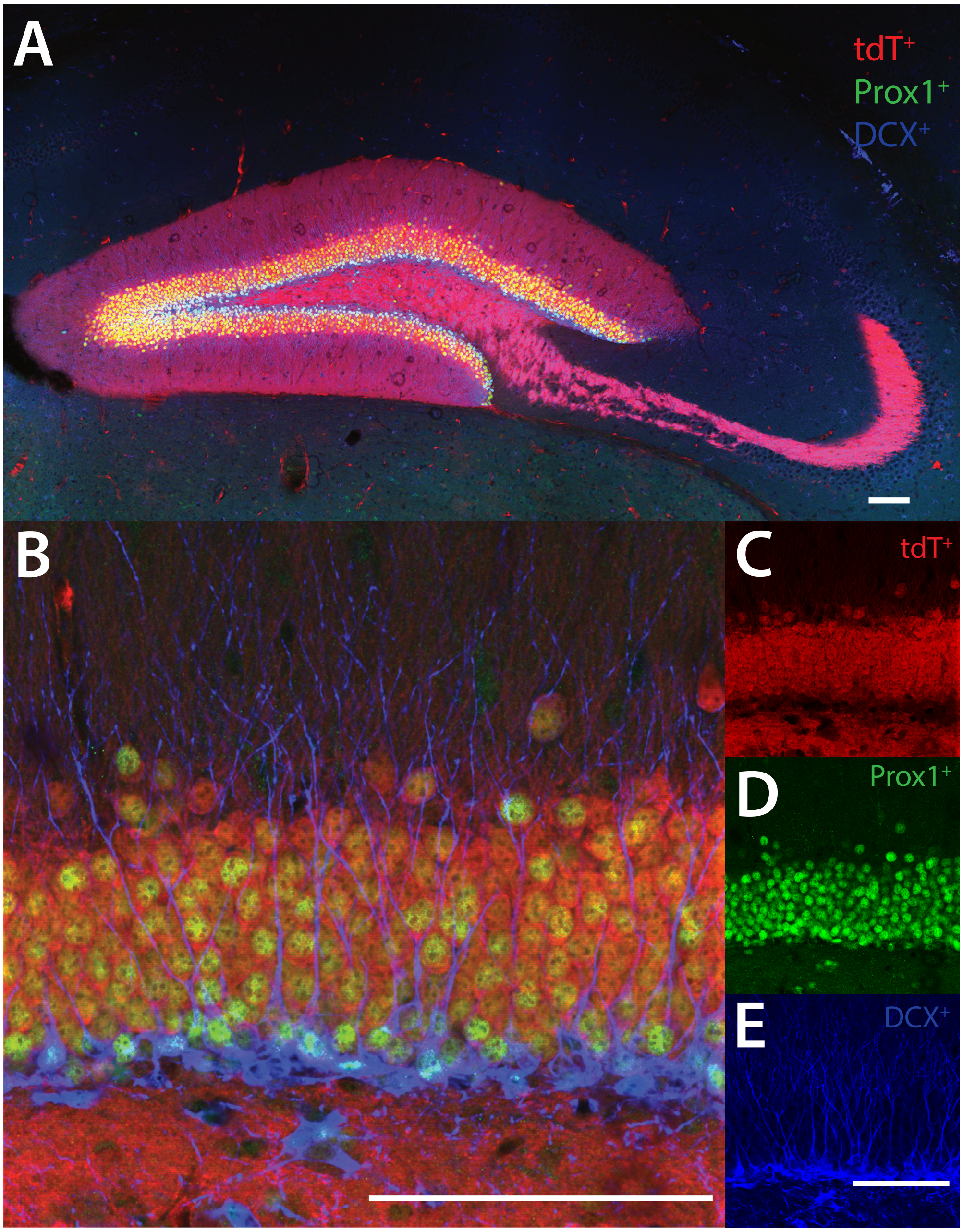
